# Intrahost speciations and host switches shaped the evolution of herpesviruses

**DOI:** 10.1101/418111

**Authors:** Anderson F. Brito, John W. Pinney

## Abstract

Cospeciation has been suggested to be the main force driving the evolution of herpesviruses, with viral species co-diverging with their hosts along more than 400 million years of evolutionary history. Recent studies, however, have been challenging this assumption, showing that other co-phylogenetic events, such as intrahost speciations and host switches play a central role on their evolution. Most of these studies, however, were performed with undated phylogenies, which may underestimate or overestimate the frequency of certain events. In this study we performed co-phylogenetic analyses using time-calibrated trees of herpesviruses and their hosts. This approach allowed us to (i) infer co-phylogenetic events over time, and (ii) integrate crucial information about continental drift and host biogeography to better understand virus-host evolution. We observed that cospeciations were in fact relatively rare events, taking place mostly after the Late Cretaceous (~100 Millions of years ago). Host switches were particularly common among alphaherpesviruses, where at least 10 transfers were detected. Among beta- and gammaherpesviruses, transfers were less frequent, with intrahost speciations followed by losses playing more prominent roles, especially from the Early Jurassic to the Early Cretaceous, when those viral lineages underwent several intrahost speciations. Our study reinforces the understanding that cospeciations are uncommon events in herpesvirus evolution. More than topological incongruences, mismatches in divergence times were the main disagreements between host and viral phylogenies. In most cases, host switches could not explain such disparities, highlighting the important role of losses and intrahost speciations in the evolution of herpesviruses.

## INTRODUCTION

Herpesviridae is a diverse family of large double-stranded DNA viruses subdivided in three subfamilies – *Alpha*-, *Beta*-, and *Gammaherpesvirinae* –, which infect different groups of vertebrates, including birds, mammals and reptiles (Davison et al. 2009). The evolutionary history of herpesviruses (HVs) dates from the Early Devonian, around at least 400 Millions of years ago (Mya) (McGeoch et al. 2006). Up to recent years, HVs were considered to be species-specific, and to have evolved alongside their hosts mainly by cospeciation (McGeoch et al. 1995; Davison 2002; Jackson 2005; McGeoch et al. 2006), an event that implies concurrent and interdependent splits of host and viral lineages over time (de Vienne et al. 2013). As more information about the divergence timing of host and viral species became available, the predominance of cospeciation as the main mechanism of HV evolution started to be challenged, especially due to mismatches between the divergence times of hosts and viruses, which evoked alternative hypothesis to explain HV evolution, such as host switches (transfers) (Davison 2002; Escalera-Zamudio et al. 2016).

Transfers take place when viruses succeed in infecting a new host still unexplored by their ancestors (de Vienne et al. 2013). Recent studies have been suggesting that host switches are probably more frequent than previously thought (Escalera-Zamudio et al. 2016; Geoghegan et al. 2017), but detecting host switches can be difficult, as extinction of viruses transmitted to new hosts may occur frequently (Geoghegan et al. 2017). Duplication of viral lineages (intrahost speciation) is another evolutionary process playing an important role on host-pathogen evolution (de Vienne et al. 2013). By means of intrahost speciations, multiple species of viruses can explore a single host species, especially when new lineages occupy distinct biological niches (tissues) within the hosts (Davison 2002). Finally, another event playing a key role in host-virus evolution is the loss of viral lineages, events that in a co-phylogenetic context can mean: (i) symbiotic extinction (Lovisolo et al. 2003); (ii) sorting events (Johnson et al. 2003); or even (iii) rare/undiscovered species, as a result of undersampling (Page and Charleston 1998).

The aforementioned mechanisms can be inferred using co-phylogenetic analyses, such as tree reconciliations, which help us understand the relationship between hosts and parasites over time (Page and Charleston 1998). To do that, such methods identify differences and similarities between the topologies of host and parasite trees, where congruence may indicate points of co-divergence (cospeciation), while incongruences my imply host switches or intrahost speciations followed by losses (de Vienne et al. 2013). Topological congruence, however, not always is caused by cospeciation events: similar topologies of host and parasite phylogenies can happen by chance, as a result of repeated host switches (de Vienne et al. 2013). To better understand the intricate evolutionary processes of HVs, co-phylogenetic analyses can be applied to elucidate the pathways taken by viruses while their hosts and the environment evolve. By integrating time-calibrated phylogenies and historical biogeographical information, in this study we present detailed scenarios unravelling the evolution of herpesviruses with their hosts, providing answers to the following questions: (*i*) What are the main co-phylogenetic events shaping HV evolution?; (*ii*) How have herpesviruses achieved their broad host range?; (*iii*) Have the three HV subfamilies evolved following similar strategies?

## RESULTS

### Phylogenetic analysis

Herpesvirus evolutionary history is older than that of their extant hosts (McGeoch et al. 2006). The MRCA (Most Recent Common Ancestors) of all hosts date back between 352 and 307 Mya, while the MRCA of all HV subfamilies existed at least between 416 and 373 Mya. As evidenced by the tanglegram in Figure 1, despite the apparent congruence of some clades in both phylogenies, topological disagreements are common for most virus-host pairs, as highlighted by the overcrossing connections between the trees.

**Figure 1.**
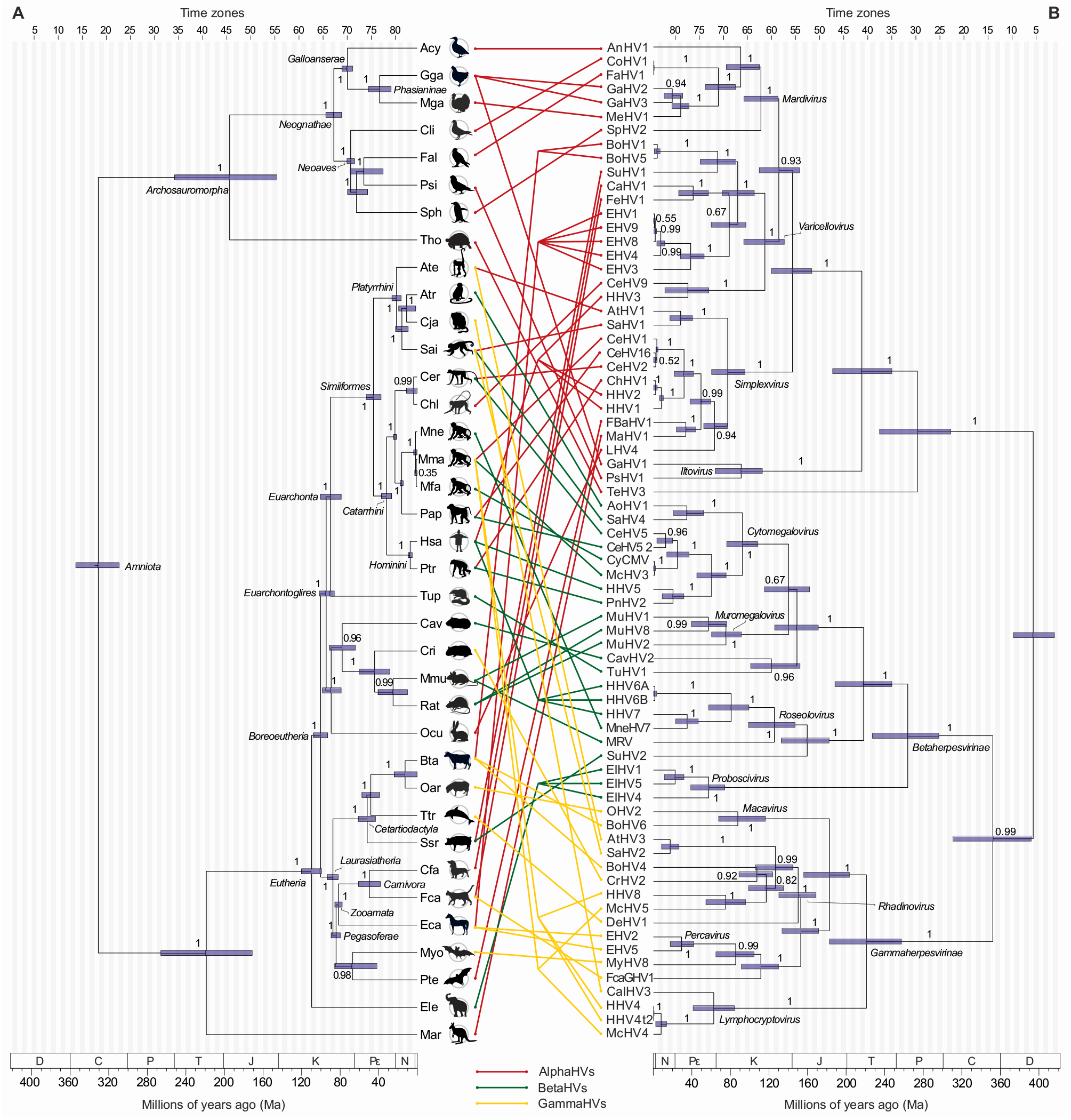
Host-virus tanglegram. Viruses (right) are connected to their hosts (left) with lines whose colours denote the three herpesviral subfamilies: *Alpha*- (red); *Beta*- (green) and *Gammaherpesvirinae* (yellow). Node height HPD intervals of hosts and viruses are shown as bars, labels are provided for some taxonomic groups, and all nodes are assigned with posterior probabilities. All internal nodes in the host tree were time-calibrated with priors. In the viral tree, priors were assigned to only two nodes (the root node and HHV1-HHV2 MRCA), while the others had their date ranges estimated from sequence data. Both trees are divided in time zones of 5 Myr, as shown by the scale at the top, and some node HPD intervals span more than one time zone. The geologic time scale is set according to (Gradstein et al. 2012), where D = Devonian period; C = Carboniferous; P = Permian; T = Triassic; J = Jurassic; K = Cretaceous; Pε = Paleogene; N = Neogene; and * = Quaternary period. Host acronyms are defined as follows: Acy = *Anser cygnoides*; Ate = *Ateles sp.*; Atr = *Aotus trivirgatus*; Bta = *Bos taurus*; Cav = *Cavia porcellus*; Cer = *Cercopithecus aethiops*; Cfa = *Canis lupus familiaris*; Chl = *Erythrocebus patas*; Cja = *Callithrix jacchus*; Cli = *Columba livia*; Cri = *Cricetidae*; Eca = *Equus caballus*; Ele = *Elephas maximus*; Fal = *Falco mexicanus*; Fca = *Felis catus*; Gga = *Gallus gallus*; Hsa = *Homo sapiens*; Mar = *Macropodidae*; Mfa = *Macaca fascicularis*; Mga = *Meleagris gallopavo*; Mma = *Macaca mulatta*; Mmu = *Mus musculus*; Mne = *Macaca nemestrina*; Myo = *Myotis velifer*; Oar = *Ovis aries*; Ocu = *Oryctolagus cuniculus*; Pap = *Papio sp.*; Psi = *Amazona oratrix*; Pte = *Pteropus sp.*; Ptr = *Pan troglodytes*; Rat = *Rattus sp.*; Sai = *Saimiri sp.*; Sph = *Spheniscus sp.*; Ssr = *Sus scrofa*; Tho = *Testudo horsfieldii*; Ttr = *Tursiops truncatus*; and Tup = *Tupaiidae*. For more details on host and viral taxonomy, accession numbers, and TaxID can be found in S1 Table.

#### Cost regimes

The co-phylogenetic analysis of 72 herpesviral species and their respective 37 hosts has revealed different possible scenarios to explain their evolution, according to distinct event costs. We tested 256 cost regimes, which favoured or penalized events differently (see supplementary tables S2-S4). Since the association of internal nodes in viral and host trees was constrained by their time zone ranges, the total number of possible solutions was limited, with many cost regimes resulting in similar solutions, which differed mostly in terms of the time zones where the inferred events probably took place. As shown in Figure 2, except for AlphaHVs (Figure 2A), the number of possible solutions (i.e. the number of events inferred for each event type, in distinct cost regimes) was very restricted in reconciliations involving BetaHVs and GammaHV, with both showing two main scenarios (Figure 2B and C): one with high numbers of Intrahost Speciations (IS) and Losses (LO), resulting in less likely solutions; and another one where such events were inferred at much lower numbers, mostly matching the median values (grey horizontal bars). For AlphaHVs, such fluctuation between high and low numbers of IS and LO was also observed, but more continuously distributed at both extremes.

**Figure 2.**
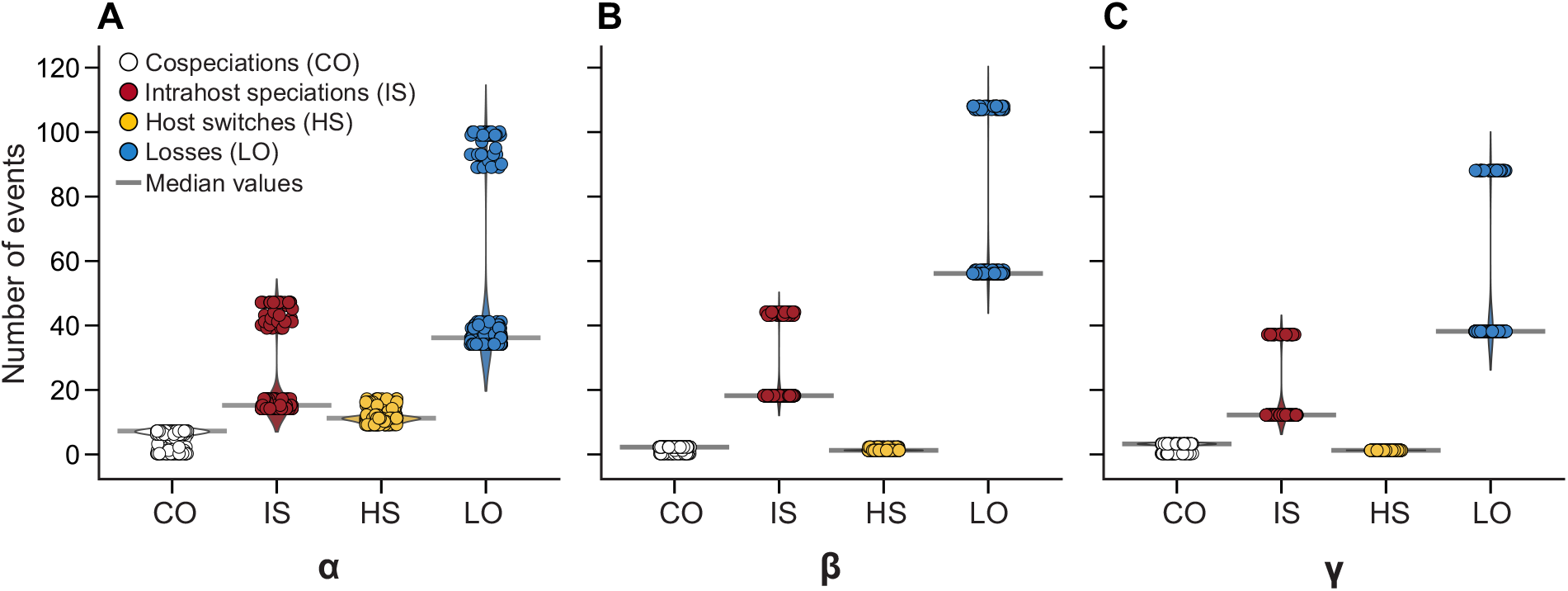
Co-phylogenetic events inferred in reconciliations between virus and host trees. A total of 256 distinct cost regimes were explored, resulting in distinct, but relatively similar scenarios explaining the disagreements between viral and host evolutionary histories. The main differences among the inferred scenarios concerned the number of Intrahost Speciations (IS, red) and Losses (LO, blue), two closely associated events. The panels show the number of inferred events under distinct cost regimes, in virus-host tree reconciliations involving A) AlphaHVs, B) BetaHVs and C) GammaHvs. Based on the median number of events for each event type (grey bars), an optimal cost regime was selected.

For each subfamily, a single cost regime was selected according to the following order of criteria, it should: (1) reconstruct the median number of events for each event type (as shown by grey bars on Figure 2); (2) ensure the highest possible support values for reconstructed events, and; (3) output the lowest possible overall cost. As a result, an optimal cost regime with the following relative costs was selected to explain the topological disagreements between viral and host tree: Cospeciation (CO) = 0; Intrahost speciation (IS) = 1; Host Switch (HS) = 2; and Loss (LO) = 0. Since the reconciliations between HVs from different subfamilies were analysed separately, it allowed us to examine the predominance of each co-phylogenetic event across different HV genera (Figure 2), as shown in the following sections.

#### Cospeciations over the last 100 millions of years

Cospeciations were only found after the Late Cretaceous (~100 Mya), and have shown to be a rare event among HVs (Figure 2 and 3). For alphaherpesviruses, seven cospeciation events were reconstructed (Table 1), the oldest one assigned to the Late Cretaceous (~92-81 Mya), involving ancestors of *Varicellovirus* infecting *Laurasiatheria* mammals (see Figure 4). This event was followed by at least other two cospeciations, one at the split of ancestors of canines, felines and equines (*Zooamata*); and another at the split of *Carnivora* ancestors. Among mardiviruses, ancestors of GaHV1 and PsHV1 (genus *Iltovirus*) co-diverged with bird ancestors (*Neognathae*) around 78-94 Mya, similar to what occurred for viruses infecting ancestors of chickens and turkeys (*Phasianinae*) around 36-26 Mya. Finally, among simplexviruses infecting *Catarrhini* (Old World Monkeys and Apes), two cospeciations were found, one taking place most likely at the Oligocene (~36-26 Mya); and the second at the Miocene (~9-5 Mya), involving viruses infecting ancestors of Humans and Chimpanzees (*Hominini*).

**Figure 3.**
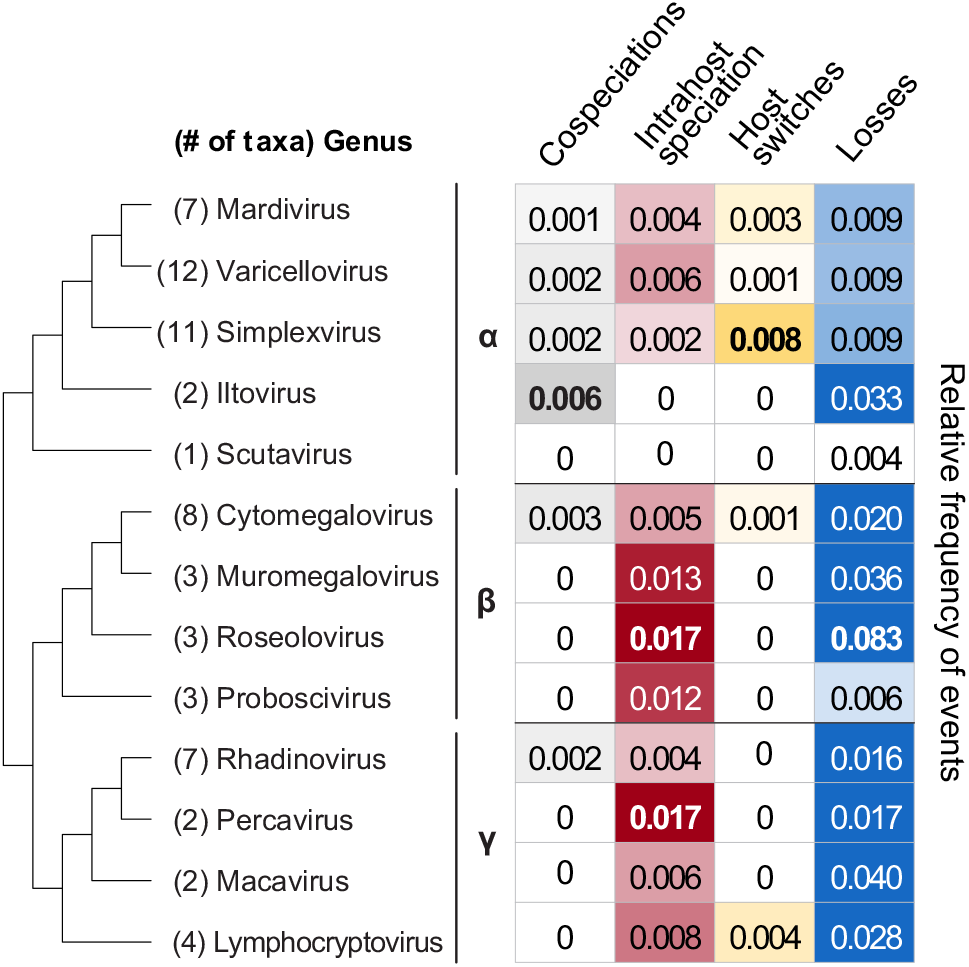
Relative frequency of co-phylogenetic events in distinct herpesviral genus, considering the optimal cost regime. Values are normalized by the total number of taxa in each genus, and their respective times to the MRCA. As highlighted, losses were the most frequent co-phylogenetic events, followed by intrahost speciations.

**Table 1.** Overall statistics of co-phylogenetic events inferred under the optimal cost regime used for virus-host tree reconciliations. The number of inferred events per HV subfamily (α, β and γ) match the median values in Figure 1. Cospeciation = (CO); Intrahost speciation = (IS); Host Switch = (HS); and Loss = (LO).

| Subfamily | Number of inferred events |  |  |  | Overall cost |
| --- | --- | --- | --- | --- | --- |
|  | CO | IS | HS | LO |  |
| $\alpha$ | 7 | 15 | 10 | 36 | 35 |
| $\beta$ | 2 | 18 | 1 | 56 | 20 |
| $\gamma$ | 3 | 12 | 1 | 38 | 14 |

**Figure 4.**
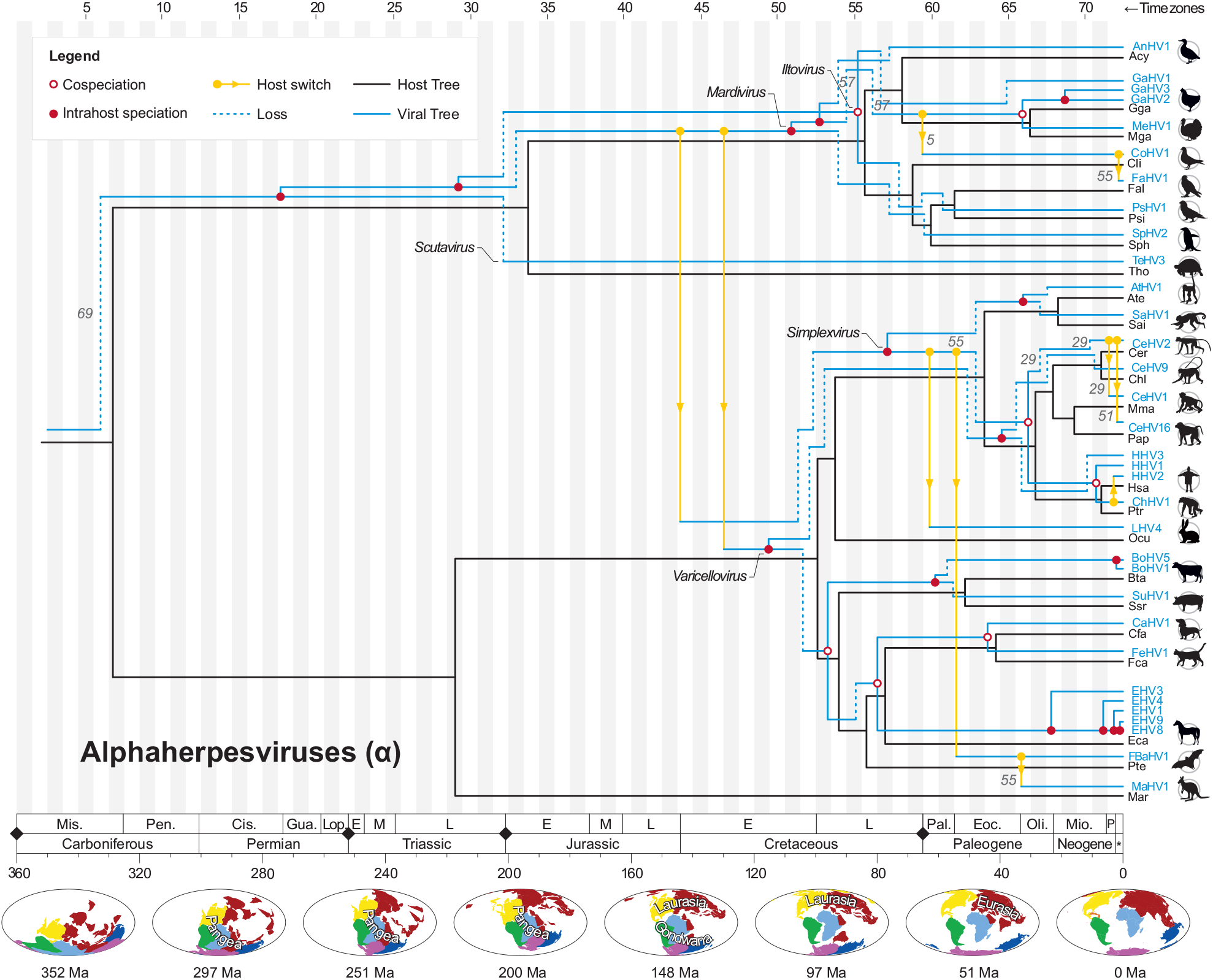
Tree reconciliation of Alphaherpesviruses and their hosts. In this representation, the host tree is shown in black, and the viral tree is shown in blue, vertically twisted, keeping its original topology and assigned time zones as shown in Figure 1. Co-phylogenetic events with support values (%) below 100 are shown. Along the time scale, black diamonds denote major events of mass extinction, as described in (Raup 1993). The maps at the bottom show changes of landmasses (continental drift) over time, and were retrieved from PBDB (Peters and McClennen 2015). Cis. = Cisuralian; E = Early; Eoc. = Eocene; Gua. = Guadalupian; L = Late; Lop. = Lopingian; M = Middle; Mio. = Miocene; Mis. = Mississippian; Oli. = Oligocene; P = Pliocene; Pal. = Paleocene; Pen. = Pennsylvanian; and * = Quaternary.

In Betaherpesviruses, only two cospeciations were reported, both along the lineage of cytomegaloviruses infecting ancestors of *Cercopithecine* (macaques and baboons) (Figure 5): the first one at the Paleogene-Neogene boundary (~24-21 Mya); and the second one around 0.8 Mya, the most recent cospeciation found in this study, involving ancestors of CyCMV and McHV3, at the split between *Macaca fascicularis* (Mfa) and *M. mulatta* (Mma).

**Figure 5.**
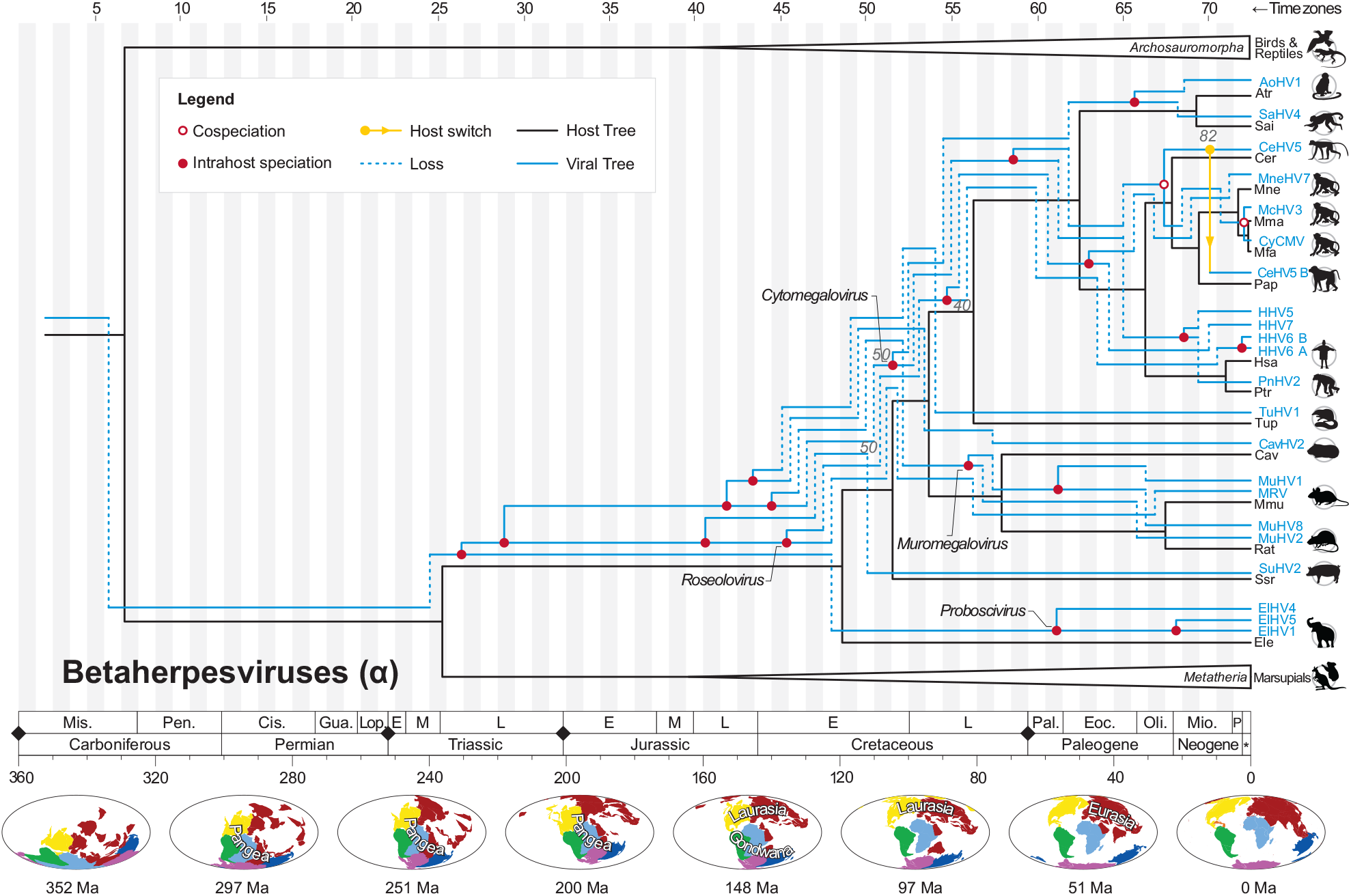
Tree reconciliation of Betaherpesviruses and their hosts.

Among Gammaherpesviruses, three cospeciations were found in different periods (Figure 6). The earliest event involved *Macavirus* ancestors, and took place at the split between *Laurasiatheria* and *Euarchontoglires* (Early Cretaceous, ~99 Mya). Following this event, HVs infecting ancestors of bats, equines and felines (*Pegasoferae*) co-diverged with their hosts at the Late Cretaceous (~88-79 Mya); and finally, at the Paleogene-Neogene boundary (~25-16 Mya), *Rhadinovirus* ancestors co-diverged with New World Monkeys (*Platyrrhini*).

**Figure 6.**
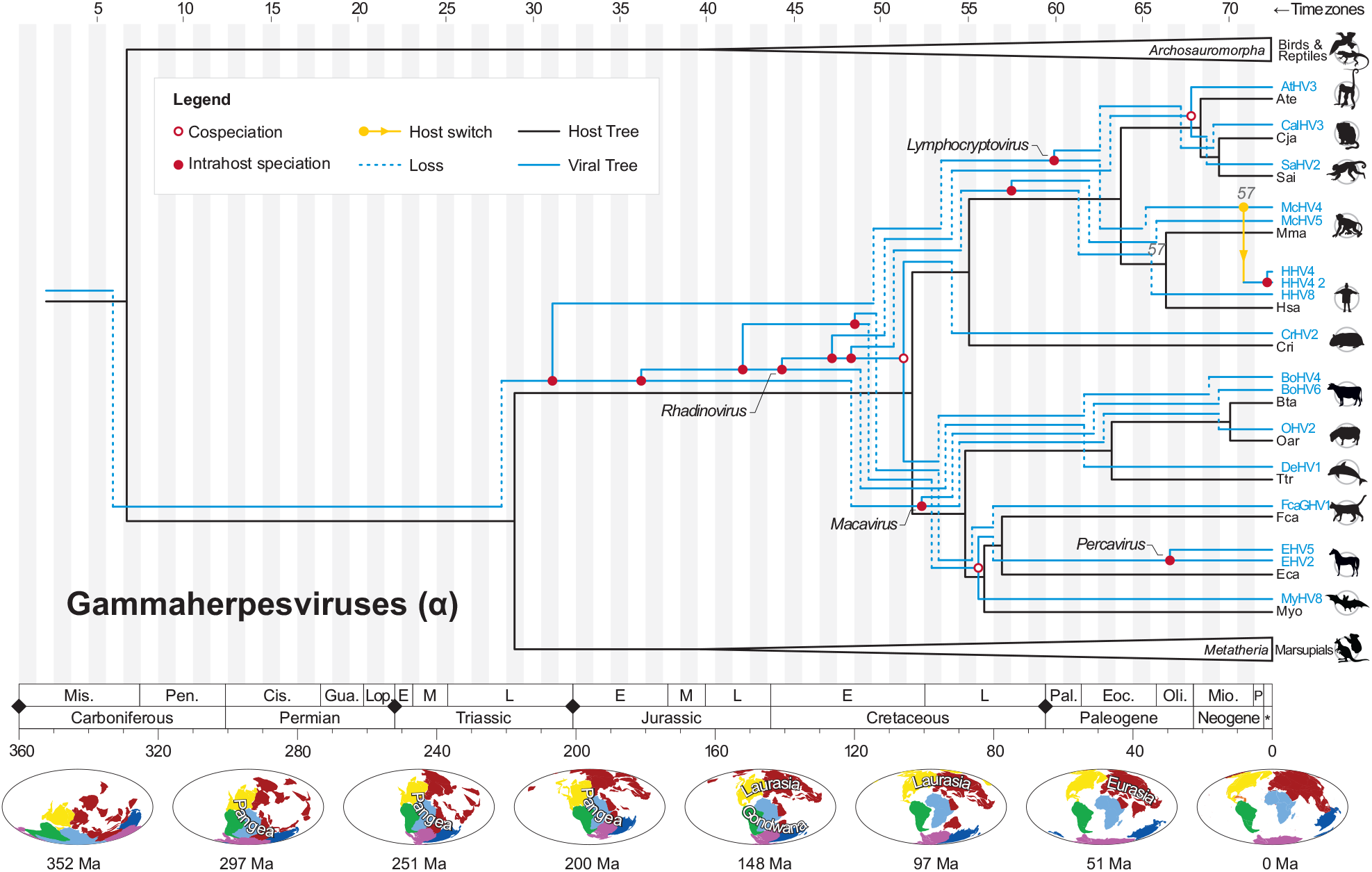
Tree reconciliation of Gammaherpesviruses and their hosts.

#### Intrahost speciations: duplications of viral lineages

Intrahost speciations (also known as duplications) occur when a parasite diverges and both lineages remain within the same host species, and represent alternative modes of evolution that explain mismatches between divergence times of viruses and hosts (de Vienne et al. 2013). Intrahost speciation was the second most frequent co-phylogenetic event in the evolution of HVs (Figure 2 and 3). Along the evolution of AlphaHVs, such events were particularly common after the Early Cretaceous. During the Permian and Triassic periods, when AlphaHVs infected ancestors of birds and reptiles (*Archosauromorpha*), two important intrahost speciations gave rise to ancestors of the main genera of *Alphaherpesvirinae* (Figure 4). Later in their evolution, along the Paleogene and Neogene, these viruses underwent a process of diversification that gave rise to multiple species sharing common hosts, as observed for Gallid (GaHV2, 3); Equid (EHV1, 3, 4, 8, 9); and Bovine alphaherpesviruses (BHV1, 5). Differently from these recent duplications, older duplication events were always followed by losses, which may explain, for example, the apparent lack of scutaviruses infecting birds, or mardiviruses and iltoviruses infecting reptiles. The evolutionary histories of BetaHVs and GammaHVs are also largely characterized by intrahost speciations (Figure 2B and 2C). As a consequence of early intrahost speciations, ancestors of BetaHV and GammaHV co-existed infecting early placental mammals from the Late Triassic to the Early Cretaceous, period when such viral lineages experienced extensive diversification (Figure 5 and Figure 6).

#### Losses: extinctions, sorting events and undiscovered herpesviruses

Among the four co-phylogenetic events, losses were the most common (Figure 2 and 3). Remarkably, considering all HV subfamilies, more than half of the losses (nearly 54%) were assigned to the Cretaceous period (~145-66 Mya). As previously mentioned, losses are usually preceded by intrahost speciations, and highlight host clades that lack viruses from certain lineages. At this point it is essential to emphasize that in the context of host-parasite tree reconciliations, losses can be interpreted in at least three distinct ways: (1) as lineage sorting events (‘missing the boat’), when a parasite fails to disperse to one of the new host species after their speciation (Johnson et al. 2003); (2) as undiscovered or rare parasites (low sampling) (Page and Charleston 1998); or (3) as genuine events of parasite extinctions (Lovisolo et al. 2003). In the latter scenario, if extinctions explain the absence of viruses infecting certain host clades, it is important to consider that points of losses in reconciliations do not reflect the exact time of the extinctions, but rather highlight a point from which such events could have happened at any subsequent time.

Throughout the evolution of herpesviruses, in Figure 4, Figure 5, and Figure 6, viral losses are depicted as dashed lines pointing towards the opposite direction the losses are assigned to. In the evolution of AlphaHVs, for example, the oldest loss dates from the Carboniferous period, before the split between sauropsids (bird/reptile ancestors) and synapsids (mammal ancestors) (Figure 4). This loss was inferred due to the potential lack of a basal group of AlphaHVs infecting synapsids in that time. Most losses of AlphaHVs were assigned to periods after the Early Cretaceous, along the earliest events of diversification of modern birds and mammals. Except for *Scutavirus*, a genus here represented by a single species (TeHV3), all genera of *Alphaherpesvirinae* show extensive losses between 110 and 70 Mya. Proportionally, losses were the most common events in the evolution of the genus *Iltovirus* (Figure 3), an old clade of HVs infecting distantly related avian families: *Psittacidae* and *Phasianidae*. Another context where losses played a central role was during the evolution of varicello- and simplexviruses infecting *Euarchontoglires* (a group that includes rabbits and primates). The evolution of these viruses alongside their hosts could only be explained by means of multiple losses, especially from ~90 Mya onwards, where at least 12 losses were observed.

Along the evolution of Beta- and Gammaherpesviruses losses were predominant. Among the BetaHV genera, *Roseolovirus* has shown the highest level of losses (Figure 3). Ancestors of these viruses originated most likely in the Early Cretaceous (~146-119 Mya), and across their evolution they were either lost in most host groups or may still exist as rare/undiscovered viruses. In GammaHVs, the genus *Macavirus* showed the second highest relative frequency of losses. Viruses in this taxonomic group are known to infect *Cetartiodactyla* hosts, mostly ruminants and swine (Ehlers et al. 2008). Since the MRCAs of these viruses and their *Laurasiatheria* hosts existed around 87 Mya (Late Cretaceous), in case they were not extinct, macaviruses infecting *Pegasoferae* (felines, canines, equines and bats) may still exist in nature.

#### Host switches

Host switches (transfers) take place when parasites succeed on infecting new hosts not yet explored by their ancestors (de Vienne et al. 2013). In all scenarios (cost regimes) investigated, host switches were invoked to explain herpesvirus-host evolution. Based on the optimal cost regime adopted in our analyses (Table 1), at least 10 host switches were reported for AlphaHVs. The oldest host switch took place when HVs from bird ancestors (sauropsids) got transferred to placental mammals in the Early Cretaceous (~144 Mya), before the split between *Mardivirus* and *Varicellovirus*. Later (~130 Mya), a second host switch took place, giving rise to ancestors of all varicelloviruses, which currently are found only in a wide range of mammalian hosts (Davison 2010) (Figure 4). Interestingly, other than these two early transfers, all the remaining host switches of AlphaHVs were assigned to periods after the Cretaceous-Paleogene boundary. In the *Mardivirus* lineage, viruses now infecting falconids and columbids (*Neoaves*) have likely originated after two host transfers: one around the Paleocene (~67 Mya) involving viruses from ancestors of modern chicken and turkeys (*Phasianinae*); and a second one in more recent times, when viruses infecting pigeons got transferred to falcons in the upper Pleistocene (~150 thousands of years ago), the most recent transfer identified in this study.

Most notably, host switches were particularly common among simplexviruses, where more than 2/3 of the transfers in AlphaHVs were observed (Figure 3). In this genus, two early transfers from primates gave rise to viruses now infecting Rabbits, Bats and Marsupials: ancestors of *Leporid alphaherpesvirus* (LHV4) switched from primates to rabbits around 76-52 Mya; and later, primate HVs switched to bats in the Eocene (~54-41 Mya), with descendants of these viruses later being transferred to marsupials in the Oligocene (~33 Mya), giving rise to the present-day *Macropodid herpesvirus* (MaHV1). The remaining transfers involved the interchange of simplexviruses among Old World Monkeys and Apes (*Catarrhini*) in the Pliocene. Ancestors of HVs now infecting monkeys of the genus *Macaca* (Rhesus, infected by CeHV1), *Papio* (Baboons, infected by CeHV16), and *Cercopithecus* (Guenons, infected by CeHV2) were likely transferred among these monkey species between 3.3 and 2.1 Mya. The exact origin of their ancestors, and the polarity of the host switches were not possible to be determined only by phylogenetic analysis, as multiple possible scenarios have shown the same overall costs (see Figure 4). Finally, an important *Simplexvirus* transfer took place between 3.6 and 0.1 Mya, when viruses infecting Chimpanzees (genus *Pan*) were transferred to humans, giving rise to HHV2, as already reported in a previous study (Wertheim et al. 2014).

In Beta- and GammaHVs, host switches were less prominent, with only one event being assigned to each of these groups, both involving primate hosts. Among BetaHVs, the species here named CeHV5-B (originally known as isolate OCOM4-52 (Blewett et al. 2015)) originated in the Miocene (~12 Mya) from viruses transferred from Guenon ancestors (genus *Cercopithecus*). Finally, for GammaHVs our results provide a possible explanation for the origins of HHV4 (Epstein-Barr Virus). Among lymphocryptoviruses, McHV4, a *Macacine gammaherpesvirus* infecting Rhesus macaques (*M. mulatta*), is the non-human HV more closely related to HHV4 (Figure 6). Ancestors of these viruses infected either Apes (early hominids) or Old World Monkeys (early cercopithecids) around 13.3 - 2.1 Mya, when these HVs switched hosts. For this particular event, the directionality of the host transfer (hominids ↔ cercopithecids) was not possible to be determined because, given the tree topologies and node heights, both directions of transfer imply the same overall cost. Despite that, other than host switch, no other co-phylogenetic event can explain the existence of these viruses in macaques and humans.

## DISCUSSION

To understand the presence of herpesviruses infecting distinct host groups, co-phylogenetic analyses with time-calibrated trees can provide consistent explanation for the dynamics of HVs evolution alongside their hosts. By using temporal data in tree reconciliations, we could also incorporate information about geological and biogeographic events that shaped life on Earth. A previous study placed the origin of herpesviruses in the Devonian (~400 Mya) (McGeoch and Gatherer 2005), a time reference used as one of the priors to calibrate the herpesviral phylogeny in this study. Based on comparative structural studies, it has been proposed that herpesviruses share a common ancestor with tailed phages (Baker et al. 2005). This hypothesis may suggest an older origin for Herpesviridae, however, the precise time range is not known, as these viruses do not produce identifiable fossil evidences. In this way, to assign divergence times to herpesvirus trees, we made use of molecular clock estimates from previous theoretical studies (McGeoch and Gatherer 2005; Wertheim et al. 2014). Despite the intrinsic uncertainties and limitations of this method, we consider such estimates to be the most objective way so far available for calibrating large DNA virus phylogenies. For this study, we calibrated the viral tree at only two nodes (the deepest node, and a shallow node), letting the remaining intermediate nodes free to have their dates estimated from sequence data. By setting constrains of monophyly for well characterized viral and host taxonomic groups, we inferred robust phylogenies with estimated ranges of divergence times, which allowed us to reconstruct, in a temporal scale, the events underlying host-viral evolution since ancient periods.

For a long time, cospeciation has been considered to be the main evolutionary mechanism driving the evolution of herpesviruses and their hosts (McGeoch et al. 1995; Davison 2002; Jackson 2005; McGeoch et al. 2006). However, most previous studies performed reconciliations using trees with few taxa, and without time scale, thus ignoring mismatches between divergence times of host and viral ancestors. For cospeciations to be accurately inferred, pathogen and host trees must not only show topological congruence, but also correspondent divergence times (de Vienne et al. 2013). Thus, the absence of temporal scale at internal nodes may overestimate the occurrence of certain events as a result of chronologically inconsistent node pairings. In the present study we performed reconciliation analyses using time-calibrated trees, which allowed us to observe that, due to temporal incompatibilities, cospeciations were in fact rare events along the herpesviral evolution, with other events playing more predominant roles.

After extensive phylogenetic characterization of herpesviruses, topological disagreements between host and viral trees became evident, and since cospeciations alone were no longer enough to explain the evolution of HVs, host transfers were initially presumed to account for such incongruences (Ehlers et al. 2008; Escalera-Zamudio et al. 2016). Contrasting with such assumption, our results revealed that while transfers played a key role on *Alphaherpesvirinae* evolution, they were uncommon among *Beta*- and *Gammaherpesvirinae*. For these two subfamilies, intrahost speciations followed by losses were abundant (Figure 2B), and may provide better explanations for the topological disagreements between their phylogenies and those of their hosts.

Viral losses do not necessarily mean extinctions, as they may also indicate undersampling or undiscovered viruses. Since the highest levels of losses were observed for herpesviral genera with few sampled taxa (Figure 3), a hypothesis of undiscovered viruses appears to be plausible, as the relative frequency of losses were to some extent influenced by the number of sampled taxa. If extinctions are invoked to explain these losses, some events linked to the evolution of the hosts can be ascribed as potential causes of the elimination of viral clades, one of them being host extinction. Although each mass extinction could have wiped out from 76% to 96% of the ancient species, most extinctions along the Phanerozoic Eon were the result of minor events taking place in-between mass extinctions (Raup 1993). Since the average duration of species is estimated to be four Myr, with genera lasting for around 28 Myr (Raup 1993), symbiont extinction may explain some of the losses inferred in the present study.

Apart from host populations being wiped out, leading to viral elimination, cataclysmic events can also cause drastic decreases in host populations (Hesse and Buckling 2016), which may promote bottleneck effects directly affecting viral adaptation to hosts, and leading to viral extinctions. It occurs especially because bottlenecks can increase the likelihood of fixation of certain host alleles, including those granting host resistance to pathogens (Hesse and Buckling 2016). Among bird species, for example, a strong bottleneck impacted avian populations after the 5^th^ mass extinction (Feduccia 2003), decreasing by nearly half the net diversification rates of birds along the Paleocene and Eocene (~66-45 Mya) (Claramunt and Cracraft 2015). Such decline in host diversity may possibly explain some of the losses inferred in clades of avian HVs, especially those belonging to *Iltovirus* (Figure 3 and Figure 4).

Since direct or indirect contact are required for viral spread into new hosts, the geological movement of landmasses can split or merge host populations, in this way allowing or preventing certain host switches (Lovisolo et al. 2003). For a better understanding of ancestral host transfers, it is important to consider the biogeography (spatial distribution) of ancestral hosts, and the geological history of the Earth. As shown at the bottom of Figure 4, Figure 5 and Figure 6, alongside the evolution of hosts and their associated viruses, the planet underwent drastic changes. All hosts included in this study are tetrapods, a group of organisms that diverged from aquatic vertebrates between 385 and 375 Mya (George and Blieck 2011). Given that the most probable tMRCA of herpesviruses dates from 416 and 373 Mya, ancestors of such viruses probably infected marine organisms, and along their terrestrialization, their viruses diverged into ancestors of the known Alpha-, Beta-, and GammaHVs. Being the first to diverge, AlphaHVs were probably lost on synapsids (proto-mammals) in the Carboniferous, reappearing as mammalian HVs much later, when avian AlphaHVs got transferred to mammals in at least two independent events (Figure 4). Such transfers, which gave rise to *Simplexvirus* and *Varicellovirus*, were already suggested in previous studies to explain the dispersal of HVs in mammalian hosts (McGeoch and Gatherer 2005; McGeoch et al. 2006). Our previous study has shown that HVs gained, duplicated and lost specific protein domains along their evolutionary history (Brito and Pinney 2020). Before and after colonizing their new hosts, simplexviruses acquired, duplicated and lost domains encoded mainly in envelope and modulatory proteins, while varicelloviruses evolved by altering their repertoire of envelope and auxiliary protein domains.

Although parasites are more likely to jump between closely related hosts, as they share similar ecological, physiological and chemical characteristics (de Vienne et al. 2013; Geoghegan et al. 2017), host switches can also occur over great phylogenetic distances, as hosts from closely related clades can independently acquire or lose immunogenetic traits (epitopes, domains or whole genes), which may change their levels of susceptibility to pathogens (Longdon et al. 2014). As host populations could have been not only physically but also genetically closer in early times than they are in current times, the origins of mammalian HVs by means of host transfers from avian ancestors could be a reasonable explanation.

In our analysis, except for the two transfers discussed above, all host switches were found in more recent times, especially after the Cretaceous/Paleogene boundary (~66 Mya). This pattern was already expected, as pointed out in a previous study, which found recent host switches as a predominant evolutionary patter in most DNA and RNA viruses (Geoghegan et al. 2017). Our results revealed that host switches were more prominent among simplexviruses. A remarkable example is the transfer involving FBaHV1 and MaHV1, which are closely related to primate HVs, but infect distantly related hosts (bats of the genus *Pteropus* and marsupials, respectively). As ancestors of *Pteropus sp.* (Megabats) inhabited Eurasia alongside primate ancestors in the Paleocene and Eocene (~59-38 Mya) (Springer et al. 2011; Springer et al. 2012), transfers of HVs from primates to bats were probably more likely to happen in that period. Finally, the subsequent host switch from megabats to macropods (Kangaroo and Wallaby ancestors) can be mainly explained by their current and ancestral distributions in Australia, and by the dispersal capabilities of bats using powered flight (Springer et al. 2011). As our previous study revealed, both FBaHV1 and MaHV1 underwent major genomic reshaping after being transferred from primates and diverge into two viral species: FBaHV1 gained and duplicated a series of envelope and modulatory protein domains, while MaHV1 lost at least eight domains, mainly present in envelope proteins (Brito and Pinney 2020). Given the defensive and offensive strategies of evolution adopted by pathogens and their hosts in molecular arms races (Daugherty and Malik 2012), herpesviruses likely succeeded at host switching by evading and/or neutralizing host immune factors by acquiring and losing elements from their domain repertoires (Brito and Pinney 2017; Brito and Pinney 2020).

Host switches involving primate HVs were observed in all herpesvirus subfamilies, most of them taking place after ~10 Mya, along the Miocene, Pliocene and Quaternary. Wertheim and colleagues (Wertheim et al. 2014) performed an in-depth analysis of the evolution of Human simplexviruses (HHV1 and HHV2) and suggested that HHV2 could have originated from host switches of HVs from Chimpanzees around 1.6 Mya. Our findings have confirmed this transfer, positioning it slightly earlier in time, around 2 Mya (95% HPD = 3.6 - 0.1 Mya), after the divergence of ChHV1 and HHV2. The same study has also pointed out another possible host switch involving ancestors of CeHV2. Our results revealed that simplexviruses infecting Old World Monkeys, such as CeHV1, CeHV2, and CeHV16 (Figure 1), underwent host switches after 4.3 Mya. Remarkably, all species of the genera *Cercopithecus* and *Papio* (except *P. hamadryas*) are found only in Africa, while those of the *Macaca* genus are mostly found in Asia (except *M. sylvanus*) (Springer et al. 2012). Ancestors of *Macaca* dispersed from Africa to Asia in the Pliocene, between 5.4 and 2.3 Mya (Springer et al. 2012), a period that matches the tMRCA of simplexviruses infecting Old World Monkeys. Thereby, our findings suggest that transfers of HVs from *Cercopithecus* to *Macaca* took place before the migration of the latter to Asia, which explains the presence of CeHV1 infecting *M. Mulatta*. Finally, between 3.5 and 0.07 Mya, transfers from *Cercopithecus* to ancestors of *Papio* took place in Africa, the place of origin and current area of distribution of most species from both genera (Springer et al. 2012).

It is known that herpesviral host switches occurred frequently in the past, especially among closely related organisms (Thiry et al. 2006; Tischer and Osterrieder 2010). As mutation rate and effective population size (*Ne*) affect the likelihood of adaptation of a pathogen in a new host (Longdon et al. 2014), the extinction of viruses transmitted to new host species may occur frequently (Geoghegan et al. 2017). As a result, most host switches cannot be easily detected, and the number of transfers inferred in our co-phylogenetic analysis is probably underestimated.

In conclusion, in this study we used time-calibrated virus-host phylogenies to perform tree reconciliations. By means of this approach we were not only able to detect topological disagreements between viral and host tree topologies, but more importantly, we reveal major chronological mismatches on divergence times of animal species and their herpesviruses, showing the relevance and fundamental contribution of temporal data for reconciling phylogenies. Our dated reconciliations highlighted the central roles of intrahost speciations in the evolution of herpesviruses, events that in most cases were followed by losses, which were predominant along the Cretaceous period. It was not possible to clarify what such losses represent in the evolution of herpesviruses, as they can represent lineage sorting events, undiscovered viruses, or even episodes of viral extinctions. As more viral samples are incorporated in tree reconciliations, the real nature of such losses will be revealed. Host switches were particularly frequent among alphaherpesviruses, especially those of the genus *Simplexvirus*. Viruses from this genus and other genera deserve further studies to uncover whether the gain or loss of specific genetic traits (domains and genes) may have favoured the colonization of new hosts or tissues (Brito and Pinney 2020). Finally, contrasting what was previously thought, cospeciations between herpesviruses and their hosts are rare events, mainly observed among alphaherpesviruses.

## MATERIALS AND METHODS

### Host/virus species and phylogenetic analysis

A total of 72 herpesviruses (subfamilies *Alpha-, Beta-, and Gammaherpesvirinae*) and their 37 host species (mammals, birds and reptiles) were included in this study (see S1 Table). Only viral species with whole genomes available on NCBI Viral Genomes Resource (Brister et al. 2015) were considered in the analysis. To infer both host and viral species phylogeny, sequences of host nuclear genes and conserved viral proteins were retrieved from NCBI. Sequence of genes encoding BDNF, CNR1, EDG1, RAG1, and RHO were used for inferring the host tree, and the viral tree was generated using sequences of orthologs of the HHV1 genes UL15, UL27 and UL30 (McGeoch et al. 2000). For host genes, when species-specific sequences were not available, sequences from related species from the same taxonomic group were used. Each gene set was aligned using MAFFT (Katoh and Standley 2013), and models of nucleotide and amino acid substitution were determined using jModelTest (Posada 2008) and ProtTest (Darriba et al. 2011), respectively. The multiple sequence alignments were used as partitions in *BEAST to infer maximum clade credibility (MCC) species trees using the Markov Chain Monte Carlo (MCMC) Bayesian approach implemented in BEAST v2.4.5 (Bouckaert et al. 2014). The analyses were performed using relaxed (uncorrelated lognormal) molecular clock, with the Yule model as a coalescent prior. The host tree was run for 500 million generations, and the viral one for 35 million generations.

The host tree had their node ages time-calibrated (in Millions of years, Myr) using as priors host divergence dates obtained from (Hedges and Kumar 2009; dos Reis et al. 2012; Claramunt and Cracraft 2015). With that, all internal nodes in the host phylogeny were assigned with lower and upper bounds for divergence times, which match the divergence dates provided at timetree.org (Kumar et al. 2017). Since fossil records or other ancestral evidences are not available for herpesviruses, viral divergence times were calibrated using theoretical estimates derived from molecular clock analyses performed by (McGeoch and Gatherer 2005; Wertheim et al. 2014). Following the same strategy used by (Brito and Pinney 2020), in the herpesvirus phylogeny, only the deepest node (root) and a shallow internal node representing the divergence between HHV1-HHV2 were assigned with time calibration priors, while the remaining nodes were left free to have had their divergence times inferred from sequence data. Constraints of monophyly were applied for viral clades based on the current taxonomic classification provided by ICTV (King et al. 2018), and host clades were constrained according to two recent and comprehensive studies (dos Reis et al. 2012; Claramunt and Cracraft 2015). Such constraints allowed the reconstruction of robust phylogenies, which fully agree with the current taxonomy of viral and host groups under study.

### Tree reconciliation

Tree reconciliations were performed separately for each HV subfamily using the program Jane 4 (Conow et al. 2010). Such analyses were performed to find low cost associations between host and viral trees, while keeping their original topologies. Taking advantage of their node height highest posterior density (HPD) intervals, all trees were converted into a Jane timed tree format using a Python script available at GitHub (see Data Availability). At this step, the continuous time scales of the phylogenies were discretized into bins of 5 Myrs (“time zones”) shared by viral and host trees. In this way, the internal nodes of host and viral trees could be assigned to specific time frames, ensuring that only nodes belonging to the same time zone could be associated for inferring potential host switches and cospeciation events, thus avoiding chronological inconsistencies. The algorithm implemented in Jane allows internal nodes to be assigned to more the one time zone, and requires that all zones should be populated with at least one host node. To meet such requirement, an outgroup clade containing artificial taxa was added to the original host tree, ensuring that their internal nodes could span time zones not originally covered by the original host nodes. A similar outgroup was added to the viral tree, allowing the pairing of artificial virus-host pairs.

Since it is not possible to ascertain the actual relative costs of events of cospeciations, intrahost speciation, host switches or losses in the evolution of HVs, we reconciled trees under multiple combinations of relative costs, in which those events were weighted with cost values varying from 0 to 3. For each HV subfamily, a total of 256 co-phylogenetic cost regimes were explored, generating several solutions, which differed in terms of overall cost and number of inferred events (see supplementary tables S2-S4). In this solution space, the median number of inferred events for each event type was calculated, and an optimal cost regime was selected based on its ability to: (i) produce the exact or similar reconciliation scenario with total number of inferred events equivalent to the median values, (ii) in which events have the highest possible supporting values, (iii) with the lowest overall cost. Following these criteria, the selected cost regime had the following relative costs of events: cospeciations = 0; intrahost speciation = 1; host switches = 2; and losses = 0 (see supplementary tables S2-S4).

## ACKNOWLEDGEMENTS

AFB is funded by *Ciência sem Fronteiras*, a scholarship programme managed by the Brazilian federal government (CAPES, Ministry of Education, Grant number: 11911-13-1). JWP is supported by a University Research Fellowship from the Royal Society. The authors thank the Imperial College London Open Access Fund for the financial support.

## DATA AVAILABILITY

All genome used mentioned in this study have their accession numbers listed as Supplementary Data. All data and codes generated in this study are deposited in the following repository on GitHub: https://github.com/andersonbrito/openData/tree/master/brito_2020_reconciliation.

## SUPPLEMENTARY DATA

Supplementary Materials are available online. Supplementary Table S1 shows the list of genomes used in this study. Supplementary Tables S2-4 lists all cost regimes used for inferring possible scenarios explaining the evolution of Alpha-, Beta, and Gammaherpesviruses.

